# Automatic recognition of macaque facial expressions for detection of affective states

**DOI:** 10.1101/2021.02.24.432760

**Authors:** Anna Morozov, Lisa Parr, Katalin Gothard, Rony Paz, Raviv Pryluk

**Author notes:** Equal contribution (Katalin Gothard, Rony Paz, Raviv Pryluk). Correspondence: R.Pa., K.G., R.Pr.

## Abstract

Internal affective states produce external manifestations such as facial expressions. In humans, the Facial Action Coding System (FACS) is widely used to objectively quantify the elemental facial action-units (AUs) that build complex facial expressions. A similar system has been developed for macaque monkeys - the Macaque Facial Action Coding System (MaqFACS); yet unlike the human counterpart, which is already partially replaced by automatic algorithms, this system still requires labor-intensive coding. Here, we developed and implemented the first prototype for automatic MaqFACS coding. We applied the approach to the analysis of behavioral and neural data recorded from freely interacting macaque monkeys. The method achieved high performance in recognition of six dominant AUs, generalizing between conspecific individuals (*Macaca mulatta*) and even between species (*Macaca fascicularis*). The study lays the foundation for fully automated detection of facial expressions in animals, which is crucial for investigating the neural substrates of social and affective states.

## Introduction

Facial expressions are a window to the internal states of an individual and a means of social communication in multiple species. The expression of emotions in man and animals was discussed first by Darwin in his eponymous treatise in which he attributed the shared features of emotional expression in multiple species to a common ancestor (Darwin 1872). Further elaboration of these ideas came from detailed understanding of the neuromuscular substrate of facial expressions, i.e., the role of each muscle in moving facial features into configurations that have social communicative value. These studies brought to light the homologies, but also the differences in how single facial muscles, or groups of muscles give rise to a relatively stereotypical repertoire of facial expressions (Ekman 1989, Ekman and Keltner 1997, Burrows, Waller et al. 2006, Vick, Waller et al. 2007, Parr, Waller et al. 2010).

The affective states that give rise to facial expressions are instantiated by distinct patterns of neural activity (Panksepp 2004) in areas of the brain that have projections to the facial motor nucleus in the pons. The axons of the motor neurons in the facial nucleus distribute to the facial musculature, including the muscles that move the pinna (Jenny and Saper 1987, Welt and Abbs 1990). Of all possible facial muscle movements, only a small set of coordinated movements give rise to unique facial configurations that correspond, with some variations, to primary affective states. Human studies of facial expressions proposed six primary affective states or “universal emotions” that were present in facial displays across cultures (Ekman and Friesen 1986, Fridlund, Ekman et al. 1987, Ekman and Friesen 1988, reviewed by Ekman, Friesen et al. 2013). The cross-cultural features of facial expressions allowed the development of an anatomically based Facial Action Coding System (FACS) (Friesen and Ekman 1978, Ekman, Friesen et al. 2002). In this system, a numerical code is assigned for each elemental facial action that is identified as an Action Unit (AU). Considering the phylogenetic continuity in the facial musculature across primate species (Burrows and Smith 2003, Burrows, Waller et al. 2006, Burrows, Waller et al. 2009, Parr, Waller et al. 2010), a natural extension of human FACS was the homologous MaqFACS (Parr, Waller et al. 2010), developed for coding the facial action units in Rhesus macaques ((for multi-species FACS review see: Waller, Julle-Daniere et al. 2020)).

The manual scoring of action units (AUs) requires lengthy training and a meticulous certification process for FACS coders, and it is a very time-consuming procedure. Therefore, considerable effort has been made towards the development of automatic measurement of human facial behaviour (Sariyanidi, Gunes et al. 2015, reviewed by Barrett, Adolphs et al. 2019). These advances do not translate seamlessly to macaque monkeys, and importantly, similar developments are desirable because macaques are commonly used to investigate and understand the neural underpinnings of communication via facial expressions (Livneh, Resnik et al. 2012, Pryluk, Shohat et al. 2020). We therefore aimed to develop and test an automatic system to classify AUs in macaques, one that would allow comparison of elicited facial expressions and neural responses at similar temporal resolutions.

Like humans, macaque monkeys do not normally activate a full set of action units required for a classical stereotypical expression, and partial sets of uncommon combination of action units are also probable and give rise to mixed or ambiguous facial expressions (Chevalier-Skolnikoff 1973, Ekman and Friesen 1976). Therefore, we chose to classify not only the fully developed facial expressions (Blumrosen, Hawellek et al. 2017) but also action units that were shown to play a role in exhibition of affective states and social communication among macaque monkeys. We test the automatic recognition of facial configurations and show that it generalizes to new situations, between conspecific individuals, and even across macaque species. Taken together, this work demonstrates concurrent validity with manual MaqFACS coding and supports the usage of automated MaqFACS in social- and affective-neuroscience research, as well as in monitoring animal health and welfare.

## Results

### The Rhesus Macaque Facial Action Coding System (MaqFACS)

There are several stereotypical facial expressions that macaques produce (Fig. 1A), that represent, as in humans, only a subset of the full repertoire of all the possible facial movements. For example, (Fig. 1B) represents three common facial expressions from the Fascicularis monkeys dataset (FD) (left, blue) and two other facial configurations that, among others, occurred in our experiments (right, yellow). Therefore, to allow the potential identification of any possible facial configuration, we chose to work in the MaqFACS domain and to recognize AUs, rather than searching for predefined stereotypical facial expressions. MaqFACS contains three main groups of AUs based on facial sectors: upper face, lower face and ears (Parr, Waller et al. 2010). Each facial expression is instantiated by a select combination of AUs (Fig. 1C).

**Figure 1.**
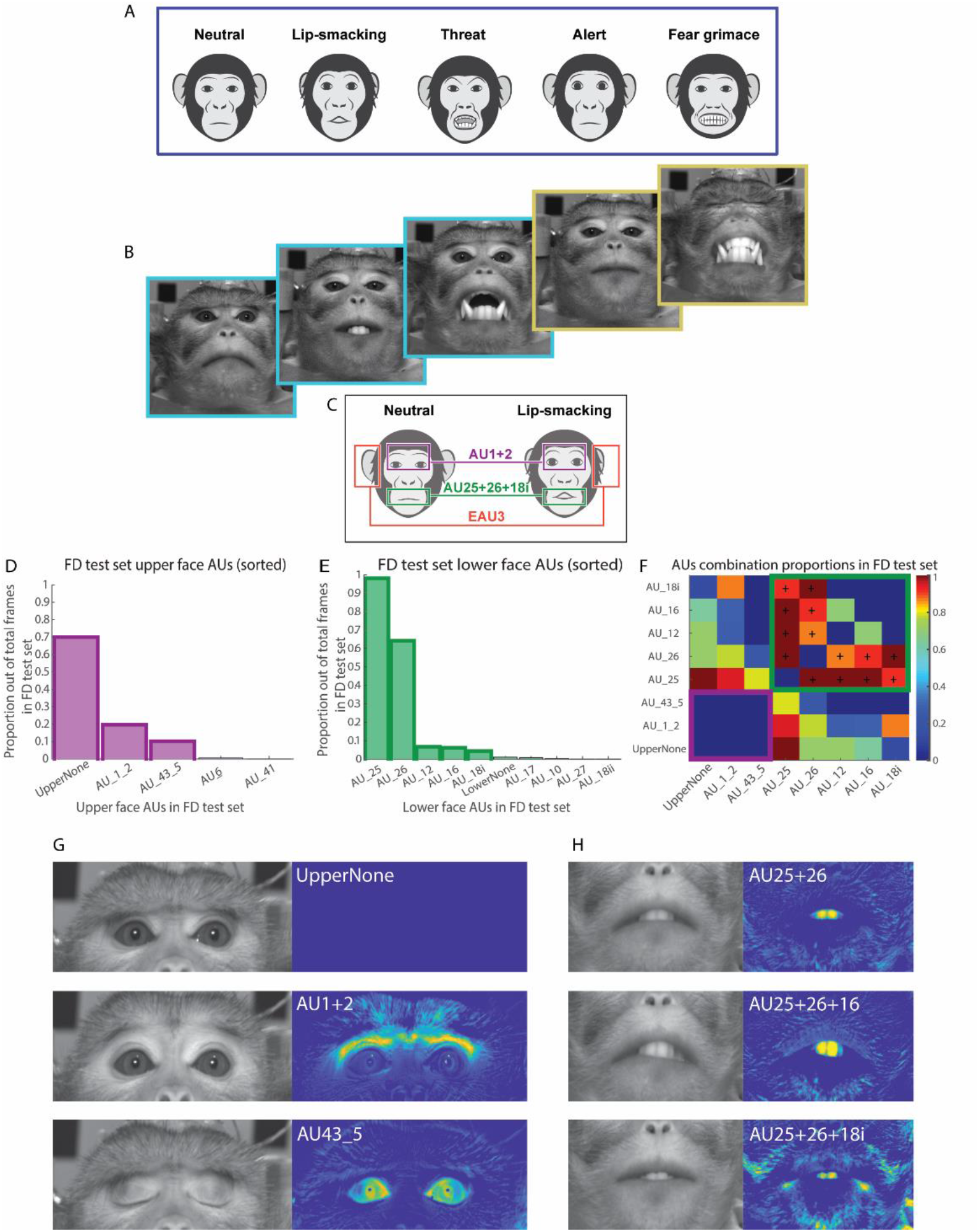
Motivation for using automatic MaqFACS to analyze facial expressions. A. The stereotypical facial expressions in macaque monkeys include the ‘neutral’, ‘lip-smacking’, ‘threat’, ‘alert’ and ‘fear grimace’ expressions (Altmann 1962, Hinde and Rowell 1962). B. Some of the facial expressions that monkeys produce during the experiments that require head immobilization match the stereotypical expressions produced during natural behaviors (for example, see the three images with blue frames on the left, correspond to the neutral, lip-smacking and threat expressions). We have also observed facial expressions that were less frequently described in the literature (two images with yellow frames on the right). C. A comparison between the neutral and lip-smacking facial expression shows that the lip-smacking example contains AU1+2 (Brow Raiser) in the upper face, AU25+26+18i (Lips part, Jaw drop and True Pucker) in the lower face, and EAU3 (Ear Flattener) in the ear region. D. The proportion of each upper face AU in the Fascicularis data (FD) test set. Bars with the solid outline (first three highest bars) represent the most frequent AUs, which were chosen for the analysis in this work. E. Same as (D) but for lower face. First five most frequent AUs were chosen for the analysis. F. Proportion matrix of AU combinations in the FD test set, for the most frequent AUs. Cells inside the magenta (bottom left) and green frames (top right) represent the combinations of upper face and lower face AUs, correspondingly. AUs that frequently occurred in combination with other AUs (in the upper face or the lower face, separately) are denoted by “+”. Cell values were calculated as the ratio between the number of frames containing the combination of the two AUs and the total frames number containing the less frequent AU. G. Left: images of upper face AUs from the FD test set. UpperNone: no coded action in the upper face. AU1+2: Brow raiser. AU43_5: Eye closure. Right: the difference of the images from the neutral face image. H. Same as (G) but for lower face. AU25+26: Lips part and Jaw drop. AU25+26+16: Lips part, Jaw drop and Lower lip depressor. AU25+26+18i: Lips part, Jaw drop and True Pucker.

The criteria for AU selection for the analysis were their frequencies and importance for affective communication (Fig. 1D,E and Materials and Methods) (Parr, Waller et al. 2010, Ballesta, Mosher et al. 2016). Frequent combinations of lower face AUs together with upper face AUs (Fig. 1F outside the magenta and green frames) may hint at the most recurring facial expressions in the test set. For example, UpperNone AU together with lower face AU25, generate a near-neutral facial expression. Considering that our aim is to recognize single AUs (as opposed to complete predefined facial expressions), lower face and upper face AUs were not merged into single analysis units. This approach is also supported by the MaqFACS coding process, which is performed separately for the lower and upper face.

Overall, our system was trained to classify 6 units: AU1+2, AU43_5 and UpperNone in the upper face, and AU25+26, AU25+26+16 and AU25+26+18i in the lower face (Fig. 1G and 1H, left and Materials and Methods). Even though AU12 was one of the most prevalent AUs in the FD test set and often occurred in combination with other lower face AUs, it was eliminated from further analysis because it appeared too infrequently in the Rhesus monkeys dataset (RD).

### Dimensionality reduction, feature extraction and classification

For each video from both datasets, seven landmark points (two corners of each eye, two corners of the mouth and the mouth center) were manually located on the mean image of frames with neutral expression. Affine transformations (geometric transformations that preserve lines and parallelism, e.g. rotation) were applied to all frames of all videos so that the landmark points were mapped to predefined reference locations (Fig. 2A, Materials and Methods). After the alignment procedure, total average image of all mean neutral expression frames was calculated. Two rectangular ROIs (regions of interest), one for the upper face and one for lower face, were marked manually on the total average image (Fig. 2B). Finally, all the frames were cropped according to the ROI windows (Fig. 2C), resulting in 396×177 pixel upper face images and 354×231 pixel lower face images. After this step, the originally RGB images were converted to grayscale. For each video, one “optimal” neutral expression frame was selected out of all the neutral expression images. Difference images (δ-images) were generated by subtraction of the optimal neutral frame from all the frames of the video (Fig. 2D, Fig. 1G and 1H, right). The main idea behind this operation was to eliminate variability due to texture differences in appearance (e.g. illumination changes), and to analyze variability of facial distortions (e.g. action units) and individual differences in facial distortion (Bartlett, Viola et al. 1996). In the last preprocessing step, upper face and lower face databases (DBs) were created by converting the δ-images to single dimension vectors and storing them as a 2-dimnesional matrix containing the pixel brightness values (one dimension is of size of the total image pixels and the second represents the images quantity). The DBs were then used for construction of training and test sets (Fig. 2E).

**Figure 2.**
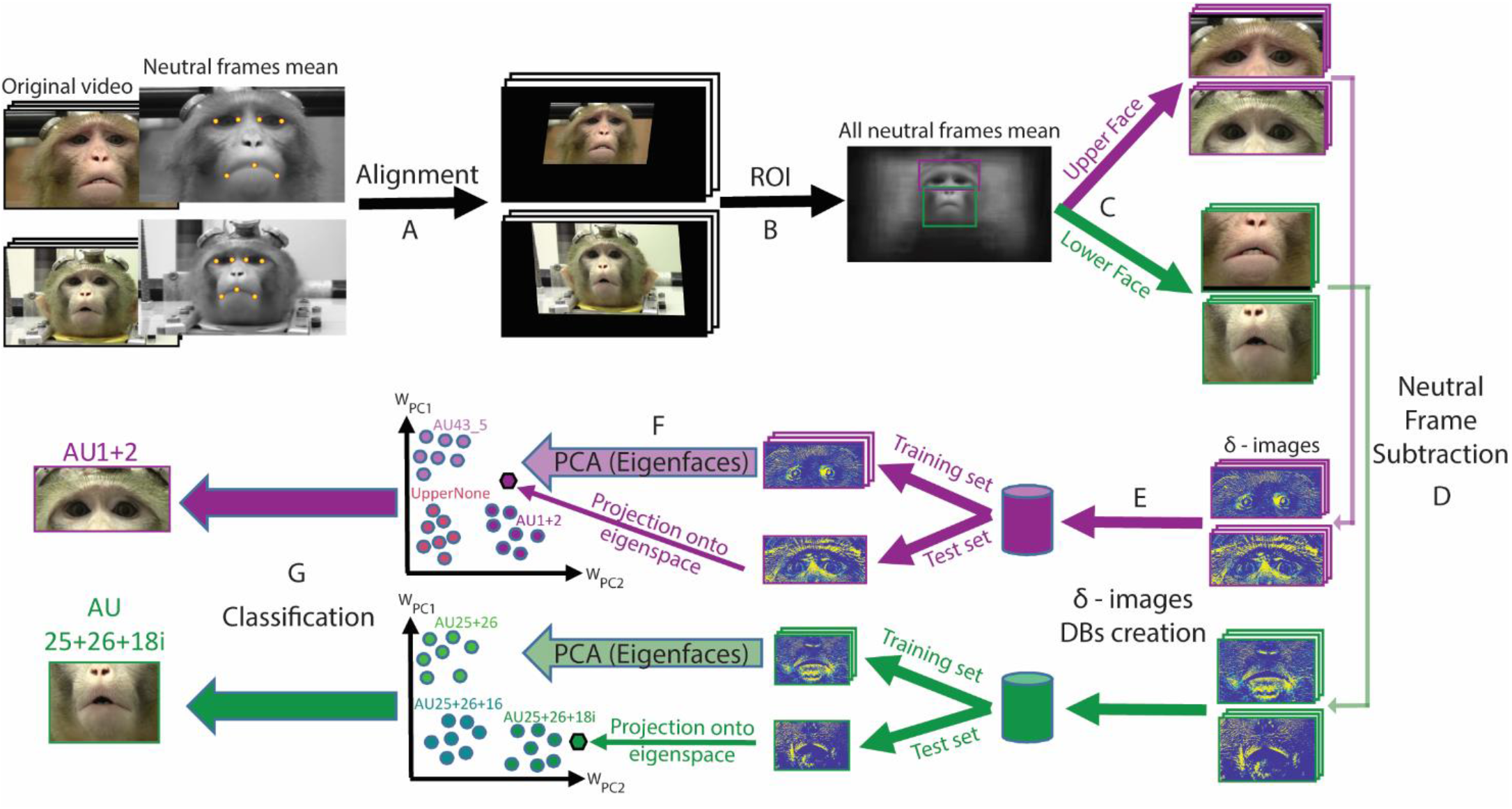
Diagram of the automatic MaqFACS AUs recognition system pipeline. A. Alignment of frames from the original video stream (example of two videos from two different Rhesus Dataset (RD) monkeys. Seven landmark points were manually selected on the mean of all neutral frames of each video. In the next step, these points were mapped to corresponding predefined positions (reference landmarks, common for all videos). The resulting affine transformation for each video was then applied to all its frames. B. Manual definition of upper face and lower face ROIs on the mean of all neutral frames. Magenta: upper face ROI, green: lower face ROI. The “All neutral frames mean” image in this scheme was calculated from all RD videos. C. Cropping of all the frames according to upper face and lower face ROIs. D. Generation of δ-images by subtracting the optimal neutral frame of each video from all its frames. The contrast and the color map of the gray scale images were adjusted for a better representation. E. Construction of lower face and upper face δ-images databases, consisting of 2-dimensional matrices where each row corresponds to one image. F. Eigenfaces extraction from the training images and projection of the training and test images onto the eigenspace (following the desired training and test sets construction). *W_PC1_* and *W_PC2_* denote the weights of PC1 and PC2, correspondingly. G. Classification of the testing images to upper face and lower face AUs. KNN (and SVM) classification was applied based on the distances between the testing and the training images in the eigenspace.

Finally, the eigenfaces analysis was applied on the training frames (the δ-images), which were first zero-meaned (Fig. 2F, Materials and Methods). Once the eigenvectors were calculated, they were normalized to unit length, and the vectors corresponding to the smallest eigenvalues (under 10^-6^) were eliminated.

To train a classification model for AUs recognition (Materials and Methods), we used the weights of the principal components (PCs) as predictors. To predict the AU of a new probe image, the probe should be projected onto the eigenspace to estimate its weights (Fig. 2F). Once the weights are known, AU classification may be applied. The output of the classifier of each facial ROI is the AU that is present in the frame (Fig. 2G). To increase the generality of our approach and to validate our results, we used both K-Nearest Neighbors (KNN) and Support Vector Machine (SVM) classifiers.

### Eigenfaces - unraveling the hidden space of facial expressions

Intuitively, light and dark pixels in the *eigenfaces* (Fig. 3A,B) reveal the variation of facial features across the dataset. To further interpret their putative meaning, we varied the eigenface weights to demonstrate their range in the training set, producing an image sequence for each PC (Fig. 3C,D). This suggests that PC1 of this upper face set (Fig. 3C top, left-to-right) codes brows raising (AU1+2) and eyes opening (AU43_5). In contrast, PC2 resembles eyes closure (Fig. 3C bottom, bottom-up). Similarly, PC1 of the lower face set (Fig. 3D top, left-to-right) probably describes nose and jaw movement. Finally, PC2 for the lower face (Fig. 3D bottom, bottom-up) plausibly correspond to nose, jaw and lip movements, reminding the transition from pushed forward lips (AU25+26+18i) to depressed lower lip (AU25+26+16).

**Figure 3:**
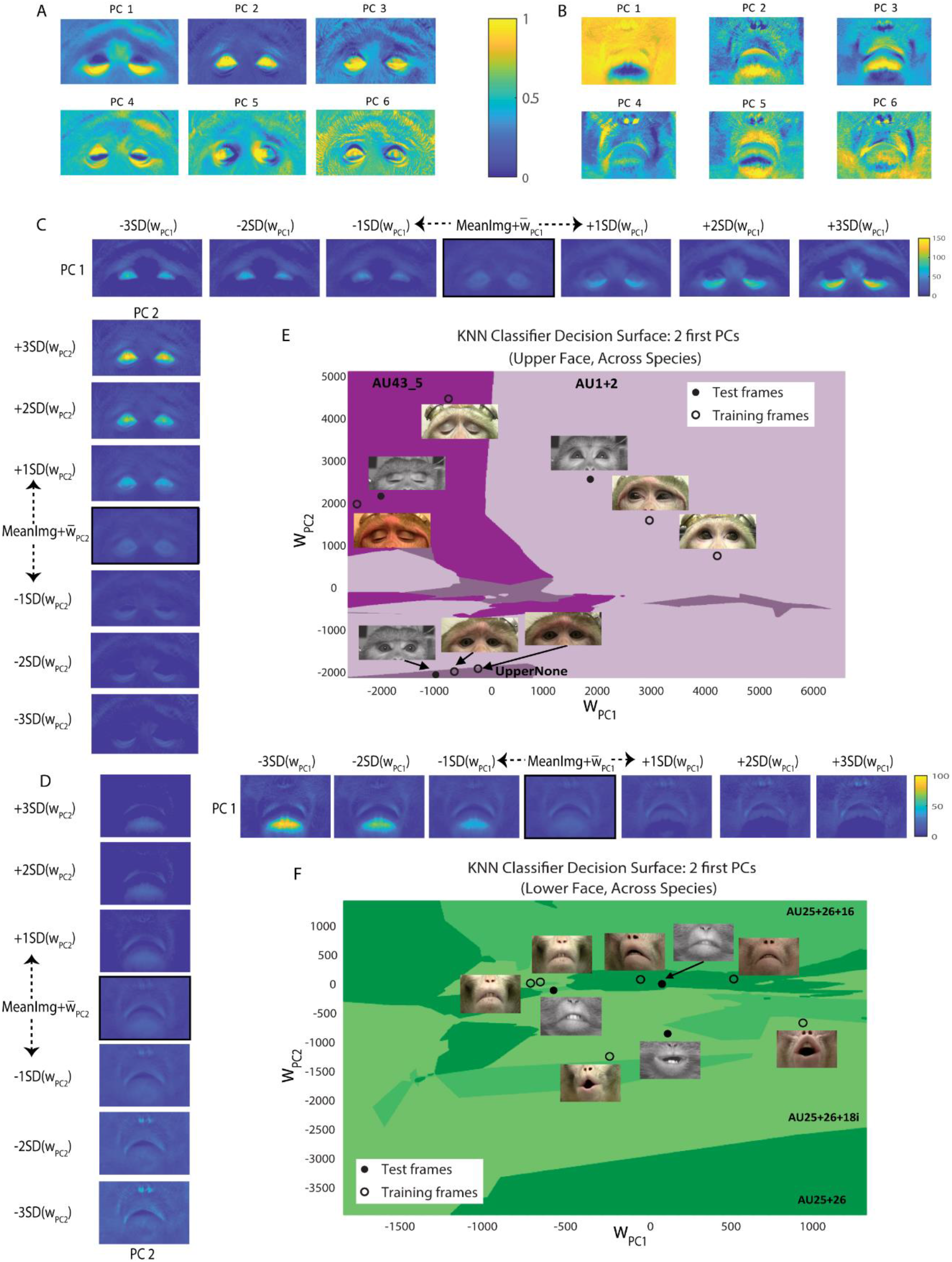
Eigenfaces analysis. A. Example of eigenfaces: six first eigenfaces (PCs) of one of the upper face training sets, containing all five Rhesus subjects from RD. The grayscale values were normalized to 0-1 range and the image contrast and color map were adjusted for a better representation. The color bar corresponds to pixel grayscale values. B. Same as (A) but for lower face. C. Example of the information coded by the first two eigenfaces. Top: the image sequence demonstrates the first eigenface from (A), added to the mean image (*MeanImg*) and varied. The middle image is the mean image of the training set (described in A), with the first eigenface added after being weighted by its mean weight (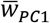). In each sequence, the weights were varied from - 3SD to +3SD from the mean weight, and the weighted PC was then added to the mean image of the training set. This procedure resulted in a different facial image for each 1SD step. The images in the sequence are ordered from left to right: the first image contains the variation by -3SD (i.e. PC1 weighted by -3SD of its weights and added to the middle image), and the last one is the variation by +3SD. Bottom: same as top but for the second eigenface (PC2). The image sequence is ordered from bottom to top. The grayscale values were normalized to 0-150 range and the image contrast and color map were adjusted for a better representation. The color bar corresponds to pixel grayscale values, and is mutual for both top and bottom schemes. D. Same as (C) but for lower face and with grayscale normalization to range 0-100. E. Example of decision surface for upper face KNN classifier, trained for generalization *across species*. The training set is the one described in (A) and the test set is Fascicularis monkey D frames from FD. The decision surface is presented along the first two dimensions – weights of PC1 and PC2 (*w_PC1_* and *w_PC2_*, correspondingly). Each colored region denotes one of the three upper face AU classes. The frames in color are training set images and the gray-scaled ones are from the test set. The classification decision is based on the test frames’ proximity to samples of a certain class in this compressed subspace. For better illustration, the images shown here are frames after alignment, but before the neutral frame subtraction. F. Same as (E) but for the lower face and Fascicularis monkey B from FD test set.

To illustrate the *eigenspace* concept, we present decision surfaces of two trained classifiers (Fig. 3E,F), along their first two dimensions (the weights of PC1 and PC2) which account for changes in facial appearance in (Fig. 3C,D). We show several training and test samples along with their locations following the projection onto the eigenspace. The projection of the samples is performed to estimate their weights, which are then used by the classifier as predictors.

### Parameter selection

Example of parameter selection (Materials and Methods) for a Fascicularis subject is shown in Fig. 4A. Interestingly, this upper face classification required much larger pcExplVar (93% versus 60% in the lower face; the difference observed in both Fascicularis subjects). Specifically, this upper face classifier achieved its best performance with 264 PCs, opposed to the lower face classifier succeeding with only 15 PCs (Fig. 4B). The most likely explanation is the large difference between the training-set sizes (3639 upper face versus 930 lower face images). Additionally, the eye-movement in the upper face images may require many PCs to express its variance.

**Figure 4:**
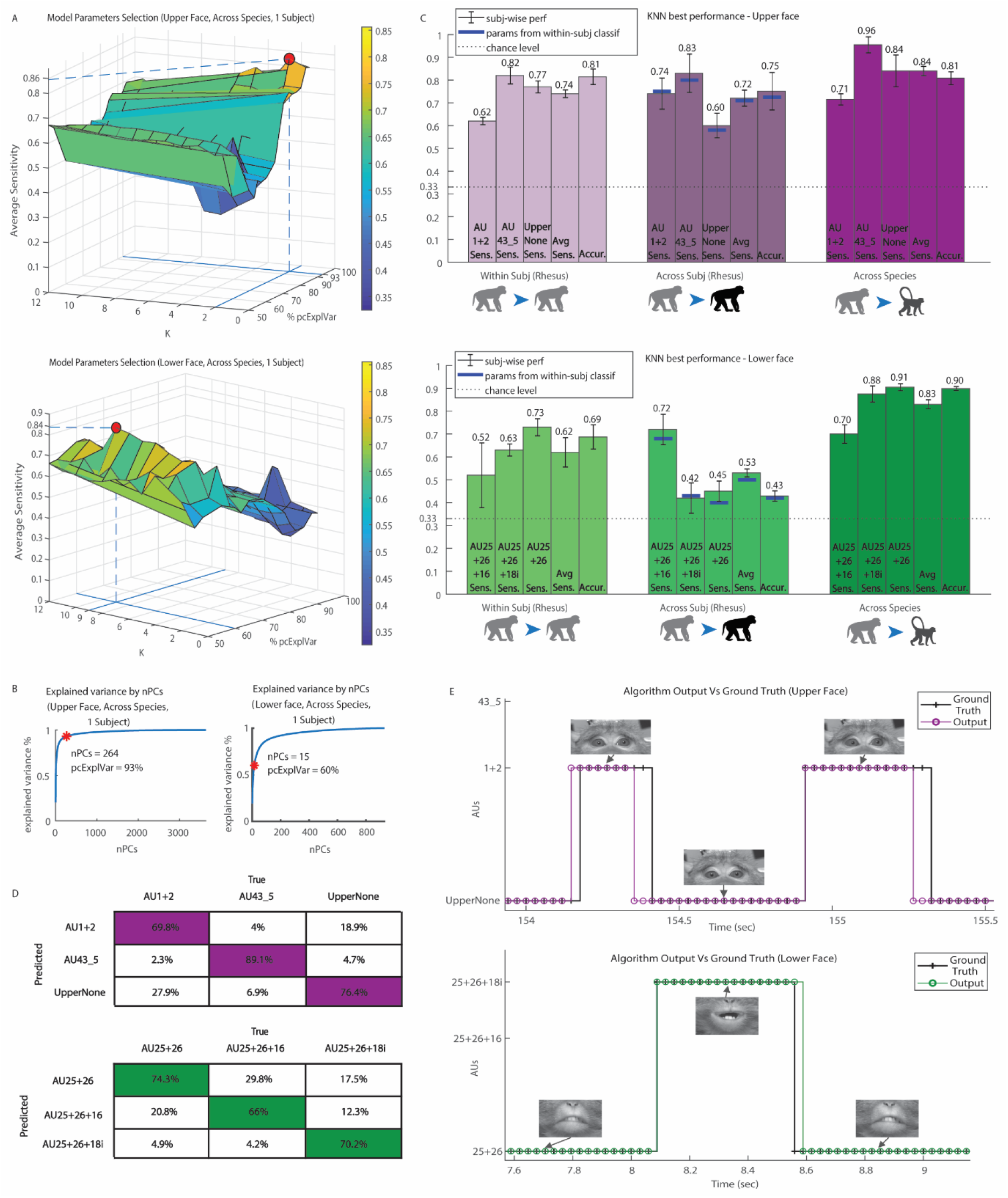
Results - parameters selection and model performance. A. Top: example of parameter selection for upper face KNN classifier, trained for generalization *across species*. The training set in the example is the one described in fig. 3(A), the test set is monkey D frames from FD and the distance metric is set to be Euclidean. The surface represents the performance of KNN classifiers with two parameters varied: k (number of nearest neighbors, varied from 1 to 12), and the percentage of the training set variance explained by the eigenfaces (“pcExplVar”, varied from 50% to 95%). Z-axis is the average sensitivity value of each model (i.e. average of the sensitivity values for the classification of three upper face AUs). The red dot denotes the highest point on the surface and hence the parameters yielding the best performance. With the selected parameters k=2 and pcExplVar = 93% the model average sensitivity value is 0.86. Bottom: Same as on top but for the lower face. The training set is one of the lower face training sets, containing all five Rhesus subjects from RD, and the test set is monkey D frames from FD. The distance metric is set to be Euclidean. The selected model has the average sensitivity of 0.84 with the parameters: k=9 and pcExplVar = 60%. B. The curves demonstrate the number of the eigenfaces that should be used to cumulatively capture a given percentage of the dataset variance. The red asterisk denotes the pcExplVar parameter value selected in (A). Left: the curve corresponds to the dataset described in (A) top. To express 93% of the dataset variance, at least 264 vectors (eigenfaces) should span the eigenspace. Right: same as left but regarding (A) bottom. To express 60% of the dataset variance, at least 15 vectors (eigenfaces) should span the eigenspace. C. Best performance of KNN classification for each generalization type. Each bar group contains five bars (from left to right): three bars describing the classifier’s sensitivity for single AUs; sensitivity averaged for three classified AUs; and the total accuracy of the classifier. The mean and the error are calculated regarding the recognition performance on a new subject. The horizontal dashed line denotes the chance level. The first bar group demonstrates the results for generalization of the classification within the same Rhesus subject (*Within Subject* (*Rhesus*): training on videos of a subject and testing on a new video of the same subject). The second group shows the generalization performance of a classifier to new Rhesus subjects (*Across Subjects* (*Rhesus*): training on videos from several subjects and testing on videos of a new subject). The blue lines denote the performance of the classifier *across subjects* using the parameters selected in *Within Subject* (*Rhesus*) case. The third group displays the generalization performance to new Fascicularis subjects (*Across Species*: training on videos from several Rhesus subjects and testing on videos of a new Fascicularis subject). In this case, the parameters should be tuned for each Fascicularis subject, and the results are the mean performance of two parameter sets (for the two Fascicularis subjects). Top: performance for upper face. Bottom: performance for lower face. D. Averaged confusion matrices of the KNN best performance results (of the three cases presented in (C)). The columns in each matrix represent the true labels, and the rows stand for the predicted labels. Top: upper face confusion matrices. Bottom: lower face confusion matrices. E. Example of the KNN classification performance demonstrating correctly recognized frames along with some recognition errors. Each data point denotes a frame in a video. The classified AUs (magenta and green lines) are shown in comparison to the ground truth labels (the black lines). Video time is displayed in the X-axis. Sample frames of the original video stream (after alignment and ROI cropping) are shown above the lines. The video for the example is taken from FD. Top: output example for upper face video. Bottom: output example for lower face video.

In contrast, the pcExplVar parameter behaved differently for generalizations *within* and *across* Rhesus subjects: their best upper face classifiers required pcExplVar of 85%, and 83% in the lower face sets. The notable difference between the parameters of these datasets suggests that one should tune a different parameter set for each dataset. Generally, the Rhesus dataset required much larger pcExplVar to describe the lower face than the Fascicularis dataset.

### Performance analysis

Overall, the best parameter set for generalization to a new video *within subject* (*Rhesus*) using KNN (Materials and Methods), performed with 81% accuracy and 74% 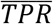 per subject for upper face, along with 69% accuracy and 62% 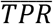 for lower face, where the chance-level is 33% (Fig. 4C, left). Best generalization *across subjects* (*Rhesus*) yielded 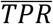 of 72% and 53% for upper and lower face respectively, with corresponding accuracy of 75% and 43% (Fig. 4C, middle), compared to 33% chance-level. The better performance in the upper face may be explained by its larger number of subjects in the CV (four in the upper face, only three in the lower face) and by greater number of examples available for training. Interestingly, applying the best parameter set of generalization *within subject* to classifiers generalizing *across subjects*, produced close-to-best performance (upper face 71% and lower face 50% 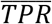). This finding suggests that tuning KNN parameters for generalization *within* Rhesus subjects, might be enough also for *across-Rhesus-subjects* generalization.

The finest results, however, were achieved in generalization *between species* with 84% 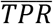 for upper face and 83% for lower face, with corresponding accuracy of 81% and 90%, concerning 33% chance-level (Fig. 4C, right). To examine whether our findings depend on the particular classification algorithm, we additionally tested this generalization with multiclass Support Vector Machine (SVM) approach. This improved the 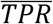 to 89% for both ROIs, indicating the advantage of using eigenfaces-based techniques for MaqFACS AUs classification.

Altogether, the upper face KNN classifiers (Fig 4D, top) separated well AU43_5, and had typical confusions between UpperNone and AU1+2. Most lower face misclassifications (Fig 4D, bottom) were between AU25+26+16 versus AU25+26 and AU25+26+18i versus AU25+26. Characteristic outputs from the system are shown in Fig. 4E.

### Behavioral analysis

To demonstrate the potential applications of our method, we used it to analyze the facial expressions produced by subject monkeys when exposed to a real life “intruder”(Fig. 5A) (Pryluk, Shohat et al. 2020). The subject monkey was sitting behind a closed shutter, when the “intruder” monkey was brought into the room (“enter” period). The shutter opened allowing the two monkeys to see each other 18 times. After the last closure of the shutter, the intruder was taken out from the room (“exit” period).

**Figure 5:**
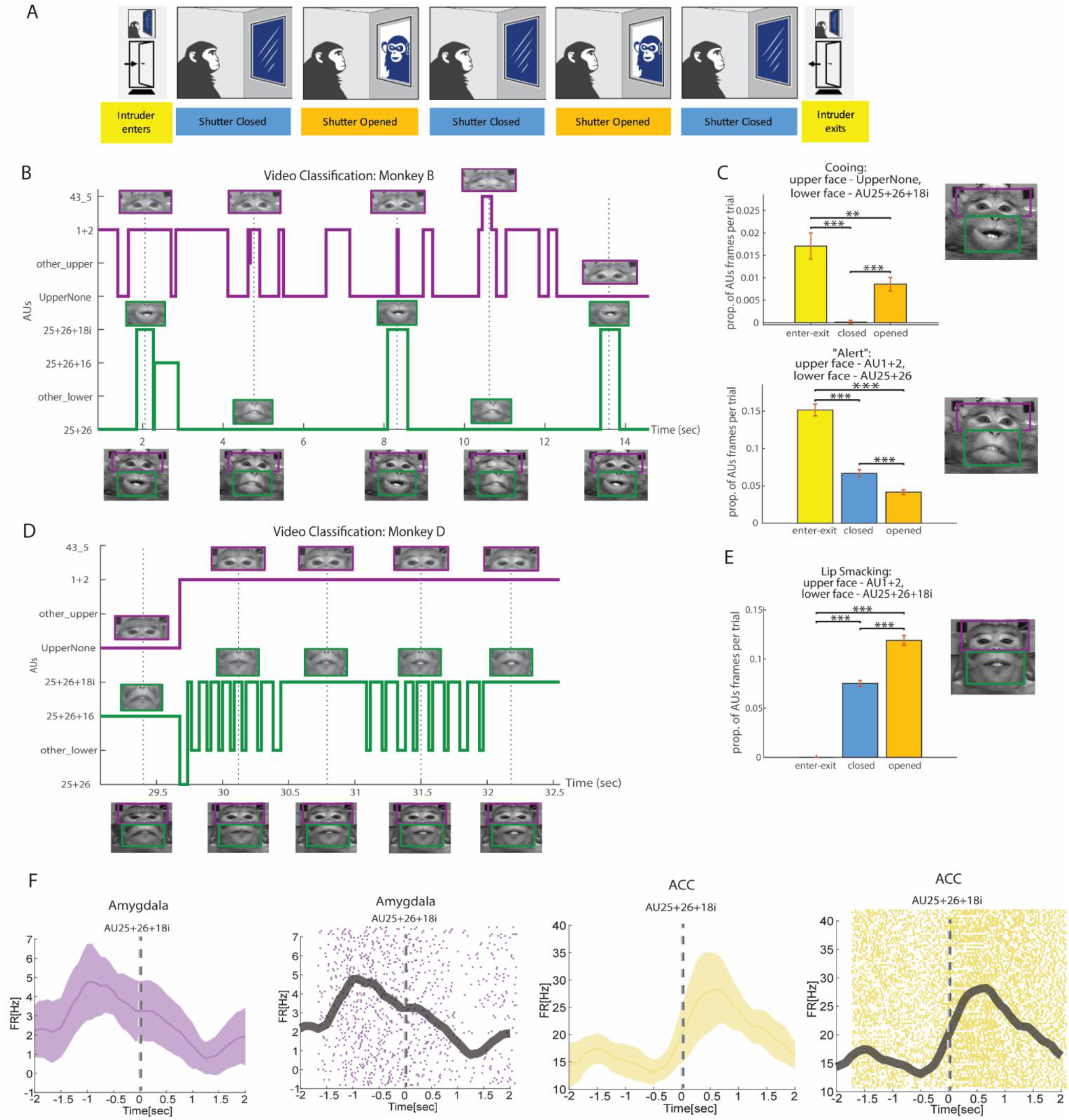
Examples of the Method Applications. A. The full behavioral paradigm is described in (Pryluk, Shohat et al. 2020). In addition, we varied the human intruder test to monkey intruder version. Monkey Intruder Block: The subject monkey sitting behind a closed shutter. The intruder monkey is brought into the room and seated behind the shutter, which remains closed. The shutter opens and closes 18 times, and the monkeys are able to see each other while it is open. The subject monkey could not see any part of the intruder unless the shutter is open. At the end of the block, the shutter closes and the intruder monkey is taken out from the room. B. Example of the final system output for monkey B from FD. Classification labels are presented on the Y-axis, while the frame time of the video-stream is on the X. “Other_upper” and “other_lower” labels are for video frames that were not part of the task of the classifier, but exist in the original video and were labeled manually. Frames of the original video (with no preprocessing) are shown on the bottom and the dashed lines denote their corresponding timing. The magenta and green lines demonstrate the outputs from the upper face and lower face algorithms, respectively. Images above the output lines exhibit the frames as they were processed in the algorithm, after alignment and ROI cropping. The estimated locations of the ROIs, comprising the full facial expressions, are illustrated in frames on the bottom by magenta and green rectangles (the positions are not precise since the original images on the bottom are not aligned). C. Facial expressions analysis following frames classification. Bars demonstrate the proportion of a specific facial configuration monkey B (from FD) elicited during one block of the experiment described in (A). This value is calculated as the ratio between frames containing the AUs combination and the total frames, per trial. Yellow bars denote the block part when the intruder monkey enters and exists the room, the blue one is for phases with the closed shutter (after the first shutter opening and before its last closure), and the orange bars stand for periods of open shutter. An example image of the analyzed expression is shown on the right (taken from the examples in B). Top: proportions of cooing facial expression events comprised of UpperNone AU for the upper face and AU25+26+18i for the lower face. Bottom: same as in top, but for “alert” facial expression – AU1+2 and AU25+25 in the upper face and lower face, correspondingly. D. Same as (B) but for monkey D from FD. E. Same as (C) but for monkey D from FD and lip-smacking facial expression with upper face AU1+2 and lower face AU25+26+18i. F. PSTHs and raster plots of one neuron in the amygdala and one in the ACC, temporally locked to the socially-associated AU25+26+18i, during monkey intruder block.

Statistical analysis of classification results for subject monkey B (Fig. 5B) revealed that in the presence of intruder, he produced several facial expressions including UpperNone and AU25+26+18i, often associated with cooing behavior. Cooing was more frequent during the “enter-exit” and open-shutter periods, than during closed-shutter periods (Fig. 5C top, χ2, p<1e-3). Moreover, subject B produced AU1+2 and AU25+26 combination more frequently during the “enter-exit” and closed-shutter periods, than during the open-shutter periods (Fig. 5C bottom, χ2, p<1e-3). We interpret this pattern as an expression of the monkey’s alertness and interest in events that were signaled by auditory but not visual inputs. Similarly, subject monkey D (Fig. 5D) produced action unit AU1+2 and AU25+26+18i together most frequently when the intruder was visible, and on occasions when the shutter was closed (intruder behind the shutter), but infrequently during the “enter-exit” periods (Fig. 5E, χ2, p<1e-3). In a social context, this pattern is associated with the lip-smacking behavior (Parr, Waller et al. 2010), representing an affiliative, appeasing social approach (Hinde and Rowell 1962).

### Neural analysis

Finally, to validate the concept and strengthen the relevancy of automatic MaqFACS for neuroscience applications, we used our method to determine whether neural activity recorded from brain regions involved in facial communication (see *Materials and Methods*) is related to specific AUs (Fig. 5A). Indeed, neurons in the amygdala and anterior cingulate cortex (ACC) were previously shown to respond with changes in firing rate during the production of facial expression (Livneh, Resnik et al. 2012). Re-analyzing the previously obtained data (Pryluk, Shohat et al. 2020) showed that neurons responded before (Fig. 5F left) or after (Fig. 5F right) the production of the socially meaningful AU25+26+18i. This finding supports the hypothesis that these regions hold neural representations for the production of single AUs or socially meaningful AU combinations.

## Discussion

This work pioneers the development of an automatic system for the recognition of facial action units in macaque monkeys. We based our approach on well-established methods that were successfully applied in human studies of facial action units (Donato, Bartlett et al. 1999). Our system achieved high accuracy and sensitivity and the results are easily interpretable in the framework of facial communication among macaques. We tested our algorithm using different macaque-videos datasets in three different configurations: within individual Rhesus monkeys, across individuals of Rhesus monkeys, and across Rhesus and Fascicularis monkeys (generalizing across species). Performance (recognition rates) was obtained for both upper and lower face and using several classification approaches, indicating that the success of this method does not depend on a particular algorithm.

We aimed to build on commonly used and well-established tools, in order to enhance applicability and ease-of-use. Interestingly, unlike the *within-Rhesus* classifications, the generalization between species required a larger number of components (explained variance) for classification of upper face AUs than for lower face AUs. This might suggest that a separate set of parameters should be fine-tuned for each dataset and ROI (lower and upper face). On the other hand, our findings show that tuning parameters for generalization *within Rhesus* subjects, might suffice also for *across-Rhesus-subjects* generalization. Further, and somewhat surprisingly, the *across-species* generalization performed better than *within-* and *across-Rhesus-subjects* generalizations. One possible explanation is that unlike in the Rhesus dataset, the Fascicularis dataset had better conditions for automatic coding, as its videos were well-controlled for angle, scale, illumination, stabilization, and occlusion. This finding has an important implication, as it shows that training on a large natural set of behaviors in less-controlled videos, can be later used for studying neural substrates of facial expressions in more controlled environments during electrophysiology (Livneh, Resnik et al. 2012, Pryluk, Shohat et al. 2020).

A direct comparison to performance of human AUs-recognition systems is not straightforward. The systems designed for humans are highly variable, due to differences in subjects, validation methods, the number of test samples and the targeted AUs (Sariyanidi, Gunes et al. 2015). In addition, some human datasets are posed, possibly exaggerating some AUs while our macaque datasets are the results of spontaneous behavior. Automatic FACS achieve great accuracy (>90%) in well-controlled conditions, where the facial view is strictly frontal and not occluded, the face is well illuminated, and AUs are posed in a controlled manner (reviewed by Barrett, Adolphs et al. 2019). When the recordings are less choreographed and the facial expressions are more spontaneous, the performance drops, (e.g. 83% in Benitez-Quiroz, Srinivasan et al. 2017). Our MaqFACS recognition system performed comparably with the human automated FACS systems despite the spontaneous nature of the macaque expressions and lack of controlled settings for the filming of Rhesus dataset.

We showed that our method can be used to add detail and depth to the analysis of neural data recorded during real-life social interactions between two macaques. This approach might pave the way toward experimental designs that capture spontaneous behaviors that may be variable across trials rather than rely on perfectly repeatable evoked responses (Krakauer, Ghazanfar et al. 2017). A departure from paradigms that dedicate less attention to the ongoing brain activity (Pryluk, Kfir et al. 2019) or internal state patterns (Mitz, Chacko et al. 2017) will increase our ability to translate experimental finding in macaques to similar finding in humans that target real-life human behavior in health and disease (Adolphs 2017). Specifically, this will allow internal emotional states and the associated neural activity that gives rise to observable behaviors to be modeled and studied across phylogeny (Anderson and Adolphs 2014). Indeed, a novel study in mice reported neural correlates of automatically-classified emotional facial expressions (Dolensek, Gehrlach et al. 2020). Finally, this system could become useful for animal-welfare assessment and monitoring (Descovich, Wathan et al. 2017, Carvalho, Gaspar et al. 2019, Descovich 2019, reviewed by McLennan, Miller et al. 2019) and in aiding the 3R framework for the refinement of experimental procedures involving all animals (Russell, Burch et al. 1959).

Given that macaques are the most commonly used non-human primate species in neuroscience, an automated system that is based on facial action units is highly desirable and will effectively complement the facial recognition systems (Loos and Ernst 2013, Freytag, Rodner et al. 2016, Crouse, Jacobs et al. 2017, Witham 2017) that address only the identity but not the behavioral state of the animal. Compared to the recently introduced method for facial expressions recognition in Rhesus macaques (Blumrosen, Hawellek et al. 2017), our system does not rely on complete stereotypical and frequent facial expressions, rather, it classifies even partial, incomplete, or ambiguous (mixed) and infrequent facial expressions, given by combination of action units. Although our system requires several manual operations, its main potential lies in automatic annotation of large datasets after tagging an example set and tuning the parameters for the relevant species or individuals. We prototyped our system on six action units in two facial regions (upper and lower face) but more advanced versions are expected to classify additional action unit combinations, spanning multiple regions of interest and tracking action units as temporal events. Further refinement of our work will likely include additional image-processing procedures, such as object tracking and segmentation, image stabilization, artifacts removal and more advanced feature extraction and classification methods. These efforts will be greatly aided by large, labeled datasets, are emerging (Murphy and Leopold 2019) to assist ongoing efforts of taking cross-species and translational neuroscience research to the next step.

## Acknowledgements

We thank Dr. Daniel Harari for comments on computer vision and machine learning techniques, and Sarit Velnchik for tagging the facial expression videos.

R.P. was supported by Israel Science Foundation Grant ISF #2352/19 and European Research Council Grant ERC-2016-CoG #724910

## Materials and Methods

### Animals and procedures

All surgical and experimental procedures were approved and conducted in accordance with the regulations of the Institute Animal Care and Use Committee (IACUC), following NIH regulations and with AAALAC accreditation.

Two male Fascicularis monkeys (*Macaca fascicularis*) and 10 Rhesus monkeys (*Macaca mulatta*) were videotaped while producing spontaneous facial movements. All monkeys were seated and head-fixed in a well-lit room during the experimental sessions.

The two monkeys produced facial behaviors in the context described in detail in (Pryluk, Shohat et al. 2020). The facial movements obtained during neural recordings have not been previously analyzed in terms of action units. Earlier experiments showed that self-executed facial movements recruit cells in the amygdala (Livneh, Resnik et al. 2012, Mosher, Zimmerman et al. 2016) and the ACC (Livneh, Resnik et al. 2012) and that neural activity in these regions is temporally locked to different socially meaningful, communicative facial movements(Livneh, Resnik et al. 2012). The video data from these monkeys was captured using two Ximea_MQ013RG (Ximea GmbH, Munster, Germany) cameras (one camera for the whole face and one dedicated to the eyes), with Kowa (Kowa Optical Products Co. Ltd., Saitama, Japan) lenses mounted on them: 16mm LM16JC10M for the face- and 25mm LM25JC5M2 for the eye-camera. The frame rates of the face- and eye-videos are 34 frames per second (~29ms) and 17 frames per second (~59ms), respectively. The size parameters are 800×700 pixels for the facial videos and 700×300 pixels for the videos of eyes. Both video types have 8-bit precision for grayscale values. The lighting in the experiment room included white LED lamps and an infrared LED light bar (MetaBright Exolight ISO-14-IRN-24, Metaphase Technologies, Philadelphia, PA, USA) for face illumination.

The 10 Rhesus monkeys were filmed during baseline sessions as well as during provocation of facial movements by exposure to a mirror and to videos of other monkeys. Videos of facial expressions of the Rhesus macaques were recorded at 30 frames per second (~33ms) rate, with 1280×720 pixels size parameters and 24-bit precision for RGB values.

### Video datasets

We used videos from two different datasets. The first, *Rhesus dataset (RD)*, consists of 53 videos from five Rhesus macaques (selected out of 10 Rhesus monkeys). Part of this dataset was used for training and testing our system within and across Rhesus subjects. The second, *Fascicularis dataset (FD)*, includes two videos from two Fascicularis macaques and was used only for testing our system across Fascicularis subjects.

All the videos in both sets capture frontal (or near-frontal) views of head-fixed monkeys. The video-frames were coded for the AUs present in each frame (none, one, or many).

The subjects and the videos for RD were selected with respect to the available data in FD, considering the scale similarity, the filming angle and the AU frequencies occurring in the videos.

### Data labeling

Video-data annotation was carried out using Noldus software “The Observer XT” (https://www.noldus.com/human-behavior-research/products/the-observer-xt). The recorded behavior coding was exported in Excel (Microsoft Excel 2016) format for further processing.

RD videos were labeled by FACS- (Friesen and Ekman 1978, Ekman, Friesen et al. 2002) and MaqFACS- (Parr, Waller et al. 2010) accredited coding expert. Another trained observer performed the coding of all FD videos according to the MaqFACS manual based on (Parr, Waller et al. 2010). Facial behavior definitions were discussed and agreed prior to the coding. To ensure consistency, we checked the inter-rater reliability (IRR) for one of the two FD videos, against additional experienced coder. Our target percentage of agreement between observers was set to 80% (Baesler and Burgoon 1987) and the IRR test resulted with 88% agreement.

All the videos were coded for MaqFACS AUs along with their frequencies and intensities. Analyzed frames with no labels were considered as frames with neutral expression. Upper- and lower-face AUs were coded separately. This partition was inspired by observations indicating that facial actions in the lower face have little influence on facial motion in the upper face and vice versa (Friesen and Ekman 1978). Moreover, neurological evidence suggests that lower and upper face are engaged differently by facial expressions and their muscles are controlled by anatomically distinct motor areas (Morecraft, Louie et al. 2001).

### AU selection

The most frequent upper face AUs in FD were the none-action AU (defined here as “UpperNone”), the Brow Raiser AU1+2 and AU43_5, which is a union of Eye Closure AU43 and Blink AU45 (Fig. 1D). The two latter AUs differ only in the movement duration, and hence were joined

There were five relatively frequent AUs in the lower face test set (Fig. 1E) that we merged into several AU groupings. All AUs that mostly co-occurred with other ones (within the same face region) were analyzed as a combination rather than single units (Fig. 1F inside the green frame). The upper face AUs however, rarely appeared as combination (Fig. 1F inside the magenta frame).

### Image preprocessing

For image height *h* and width *w*, the reference landmark points were defined by the following coordinates: (0.42w, 0.3h) and (0.48w, 0.3h) for left eye corners, (0.52w, 0.3h) and (0.58w, 0.3h) for right eye corners, (0.44w, 0.55h) for mouth left corner, (0.56w, 0.55h) for mouth right corner and (0.5w, 0.5h) for the mouth center.

### Eigenfaces: Dimensionality reduction and feature extraction

Under controlled head-pose and imaging conditions, the statistical structure of facial expressions may be efficiently captured by features extracted from Principal Component Analysis (PCA) (Calder, Burton et al. 2001). This was demonstrated in the “EigenActions” technique (Donato, Bartlett et al. 1999), where the facial actions were recognized separately for upper face and lower face images (the well-known “Eigenfaces”). According to this technique, the PCA is used to compute a set of subspace basis vectors (referred to as the ‘‘eigenfaces”) for a dataset of facial images (the training set), which are then projected into the compressed subspace. Typically, only the N eigenvectors associated with the largest eigenvalues are used to define the subspace, where N is the desired subspace dimensionality (Draper, Baek et al. 2003). Each image in the training set may be represented and reconstructed by the mean image of the set and a linear combination of its principal components (PCs). The PCs are the eigenfaces and the coefficients of the PCs in the linear combination instance their weights. The test images are matched to the training set by projecting them onto the basis vectors and finding the nearest compressed image in the subspace (the eigenspace).

### Classification

One of the benefits of the mean subtraction and the scaling to unit vectors is that this operation projects the images into a subspace where Euclidean distance is inversely proportional to correlation between the original images. Therefore, nearest neighbour matching in eigenspace establishes an efficient approximation to image correlation (Draper, Baek et al. 2003). Consequently, we employed a K-Nearest Neighbors (KNN) classifier in our system. Related to the choice of classifier, previous studies show that when PCA is used, the choice of the subspace distance-measure depends on the nature of the classification task (Draper, Baek et al. 2003). Based on this notion and other observations (Bartlett, Donato et al. 2000), we chose the Euclidian distance and the cosine of the angle between feature vectors to measure similarity. In addition, to increase the generality of our approach and to validate our results, we also tested a Support Vector Machine (SVM) classifier. To evaluate the performance of the models we define a classification trial as successful if the AU predicted by the classifier was the same as in the probe image. To further justify the classification of AUs separately for upper face and lower face ROIs, it is worth mentioning that evidence suggest that PCA-based techniques performed on full-face images lead to poorer performance in emotion recognition compared to separate PCA for the upper and lower regions (Padgett and Cottrell 1997, Bartlett 2001).

### Parameter selection

In the KNN classification, we examined the variation of three main parameters: the number of the eigenspace dimensions (PCs), the subspace distance metric and *k* - the number of nearest neighbors in the KNN classifier.

Multiple ranges of PCs were tested (the “pcExplVar” parameter), from PC quantity that cumulatively explains 50% of the variance of each training set to 95%; *k* was varied from 1 to 12 nearest neighbors and the performance was also tested with Euclidian and cosine similarity measures. For each training set and parameter set, the features were recomputed and the model performance was re-estimated. The process was repeated across all the balanced training sets (see *Data under-sampling*). The parameters of the models and the balanced training sets were selected according to the best classification performance in the validation process.

### Data under-sampling

The training sets in this study were composed of Rhesus Dataset (RD) frames from AU1+2, AU43_5 and UpperNone categories in the upper face, and AU25+26, AU25+26+16 and AU25+26+18i in the lower face (in a non-overlapping manner relatively to each ROI). For the training purposes, for both ROIs, the RD frames were randomly under-sampled 3-10 times (depending on the data volume), producing the “balanced training sets”. The main reason for this procedure was to balance the frame quantity of the different AUs in the training sets (He and Garcia 2009). For each dataset, the size of the balanced training set was defined based on the smallest category size (Table 1). As a result, for the training processes in our experiments we used upper face and lower face balanced training sets of size 3639 and 930 frames each, correspondingly.

**Table 1:**
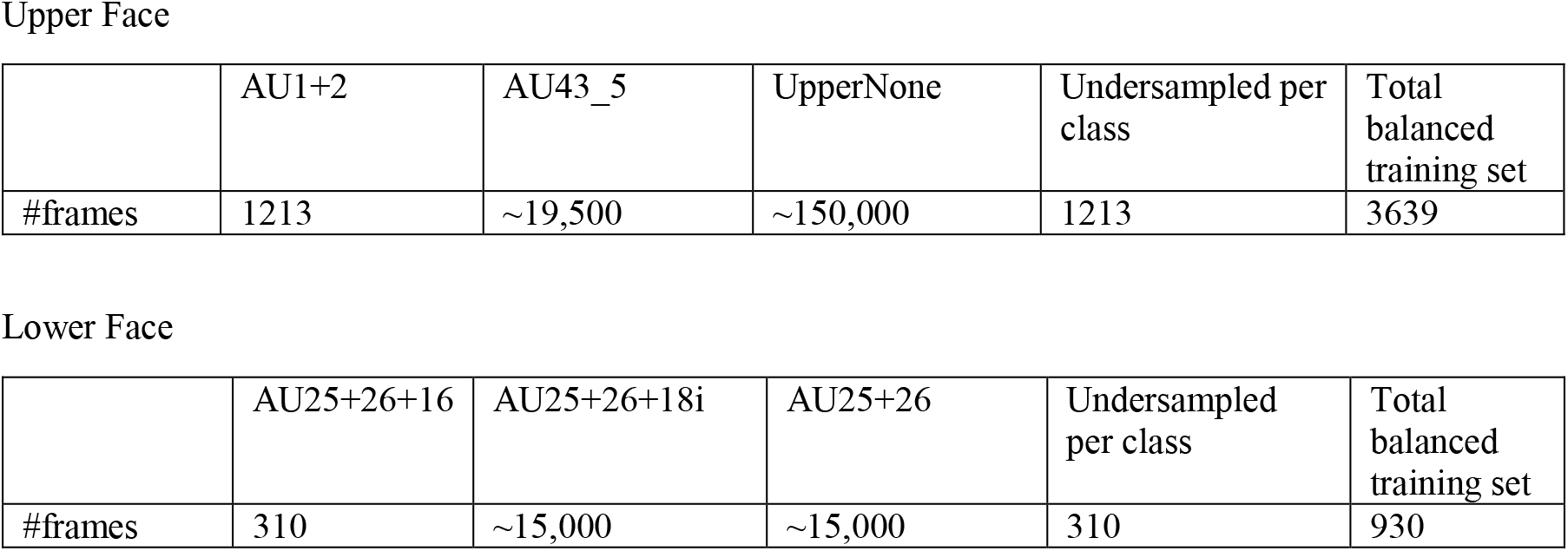
Data undersampling (RD)

It should be noted that the under-sampling procedure influences only the training but not the test sets composition (only the frames for training are selected from the balanced training sets). The test set composition depends on the subjects and the videos selected for the testing, and considers all the available frames that fit the task criteria (consequently, they are the same across all the balanced training sets).

In the upper face, the smallest category was AU1+2 with only 1213 frames (in total, from all RD subjects). On the contrary, AU43_5 category had around 19,500 frames (after eliminating RD AU45 frames due to time synchronization errors), and UpperNone class included over 150,000 images. Consequently, balanced training sets were generated each including all the AU1+2 frames, and randomly selected 1213 frames from AU43_5 along with 1213 randomly selected UpperNone frames. Therefore, the upper face balanced training sets were comprised of 3639 frames each. The same was done for the lower face, where the smallest category was AU25+26+16 with only 310 frames. Categories AU25+26+18i and AU25+26 contained over 15,000 images each. Accordingly, each lower face balanced training set included 930 frames.

### Validation and model evaluation

We tested three types of generalization. For each type of generalization, the performance was evaluated independently for upper face and lower face, using holdout validation for the Fascicularis data (Geisser 1975) and leave-one-out cross validation (CV) for the Rhesus data (Tukey 1958). The leave-one-out technique is advantageous for small datasets because it maximizes the available information for training, removing only a small amount of training data in each iteration. Applying the leave-one-out CV, data from all subjects (or videos) but one, was used for the system training, and the testing was performed on the one remaining subject (or video). We designed the CV partitions constraining equal number of frames in each class of the training sets. In both the leave-one-out CV and the holdout validation, images of the test sets were not part of the corresponding training sets, and only the training frames were retrieved from the balanced training sets. To ensure the data sufficiency for training and testing, a subject (or video) was included in the partition for CV only if it had enough frames of the three AU classes (separately for upper face and lower face).

For each generalization type, the training and the testing sets were constructed as following:

1. *Within subject* (*Rhesus*): for each CV partition, frames from all videos but one, from the same Rhesus subject, were used for training. Frames of the remaining video were used for testing. Performed on RD, on three balanced training sets. To be included in a CV partition for testing, the training and the test sets for a video had to consist of at least 20 and 5 frames per class, correspondingly. Some subjects did not meet the condition, and this elimination process resulted with three subjects for upper face and four subjects for lower face CV.
2. *Across subjects (Rhesus)*: for each CV partition, frames from all videos of all Rhesus monkeys but one, were used for training. Each test set was composed of frames from videos of the one remaining monkey. Performed on RD, on three balanced training sets. To be included for testing in the CV, the training and the test sets for a subject had to contain at least 150 and 50 frames of each class, correspondingly. In total, four subjects were included in the upper face and three in the lower face testing.
3. *Across Species*: frames from all videos of the five Rhesus monkeys were used for training. Frames from the two Fascicularis monkeys were used for validation and testing. In this case, a holdout model validation was carried out independently for each Fascicuaris monkey (each subject had a different set of model parameters selected). For this matter, each Fascicularis monkey’s dataset was randomly split 100 times in a stratified manner (so the sets will have roughly the same class proportions as in the original dataset) to create two sets: validation set with 80% of the data and test set with 20% of the data. Overall, the training sets were constructed from 10 balanced training sets of the Rhesus dataset. Validation and test sets (produced by 100 splits in total) included 80% and 20% of the Fascicularis dataset, correspondingly. The best model parameters were selected according to the mean performance in validation set (over 100 splits), and the final model evaluation was calculated based on the test set mean performance (over the 100 splits, as well).

### Performance measures

Although the balanced training sets and the CV partitions were constructed to maintain the total number of actions as even as possible, the subjects and their videos in these sets possessed different quantities of actions. In addition, while we constrained the sizes of the classes within each training set to be equal, we used the complete available data for the test sets. Since the overall classification correct rate (accuracy) may be an unreliable performance measure due to its dependency on the targets to non-targets proportion (Pantic and Bartlett 2007), we also applied a sensitivity measure (Benitez-Quiroz, Srinivasan et al. 2017) for each AU (where the target is the particular AU and the non-targets are the two remaining AUs).

We used the average sensitivity measure (average true positive rate - 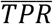) to select the best parameter set. To compare the performance of the classifiers, we present the generalization results on a *subject* (i.e., individual monkey) level (rather than *video*), for each classification type. Performance on Fascicularis dataset is reported as the mean performance of two parameter sets (one set per subject).

### Software

A custom code for automatic MaqFACS recognition and data analysis was written in Matlab R2017b (https://www.mathworks.com/). The code is available upon reasonable request.

## References

Adolphs, R. (2017). “How should neuroscience study emotions? By distinguishing emotion states, concepts, and experiences.” Social Cognitive and Affective Neuroscience 12(1): 24–31.

Altmann, S. A. (1962). “A field study of the sociobiology of rhesus monkeys, Macaca mulatta.” Annals of the New York Academy of Sciences 102(2): 338–435.

Anderson, D. J. and R. Adolphs (2014). “A framework for studying emotions across species.” Cell 157(1): 187–200.

Baesler, E. J. and J. K. Burgoon (1987). “Measurement and reliability of nonverbal behavior.” Journal of Nonverbal Behavior 11(4): 205–233.

Ballesta, S., et al. (2016). “Social determinants of eyeblinks in adult male macaques.” Scientific reports 6: 38686.

Barrett, L. F., et al. (2019). “Emotional expressions reconsidered: Challenges to inferring emotion from human facial movements.” Psychological Science in the Public Interest 20(1): 1–68.

Bartlett, M. S. (2001). Face Image Analysis by Unsupervised Learning, Kluwer Academic Publishers.

Bartlett, M. S., et al. (2000). Image representations for facial expression coding. Advances in neural information processing systems.

Bartlett, M. S., et al. (1996). Classifying facial action. Advances in neural information processing systems.

Benitez-Quiroz, C. F., et al. (2017). “Emotionet challenge: Recognition of facial expressions of emotion in the wild.” arXiv preprint arXiv:1703.01210.

Blumrosen, G., et al. (2017). Towards Automated Recognition of Facial Expressions in Animal Models. Proceedings of the IEEE International Conference on Computer Vision.

Burrows, A. M. and T. D. Smith (2003). “Muscles of facial expression in Otolemur, with a comparison to Lemuroidea.” The Anatomical Record Part A: Discoveries in Molecular, Cellular, and Evolutionary Biology: An Official Publication of the American Association of Anatomists 274(1): 827–836.

Burrows, A. M., et al. (2009). “Facial musculature in the rhesus macaque (Macaca mulatta): evolutionary and functional contexts with comparisons to chimpanzees and humans.” Journal of anatomy 215(3): 320–334.

Burrows, A. M., et al. (2006). “Muscles of facial expression in the chimpanzee (Pan troglodytes): descriptive, comparative and phylogenetic contexts.” Journal of anatomy 208(2): 153–167.

Calder, A. J., et al. (2001). “A principal component analysis of facial expressions.” Vision Research 41(9): 1179–1208.

Carvalho, C., et al. (2019). “Ethical and Scientific Pitfalls Concerning Laboratory Research with Non-Human Primates, and Possible Solutions.” Animals 9(1): 12.

Chevalier-Skolnikoff, S. (1973). “Facial expression of emotion in nonhuman primates.” Darwin and facial expression: A century of research in review: 11–89.

Crouse, D., et al. (2017). “LemurFaceID: a face recognition system to facilitate individual identification of lemurs.” BMC Zoology 2(1): 2.

Darwin, C. (1872). “The expression of emotions in men and animals.”

Descovich, K. (2019). “Opportunities for refinement in neuroscience: Indicators of wellness and post-operative pain in laboratory macaques.” ALTEX.

Descovich, K., et al. (2017). “Facial expression: An under-utilised tool for the assessment of welfare in mammals.”

Dolensek, N., et al. (2020). “Facial expressions of emotion states and their neuronal correlates in mice.” Science 368(6486): 89–94.

Donato, G., et al. (1999). “Classifying facial actions.” IEEE TRANSACTIONS ON PATTERN ANALYSIS AND MACHINE INTELLIGENCE 21(10): 974–989.

Draper, B. A., et al. (2003). “Recognizing faces with PCA and ICA.” Computer Vision and Image Understanding 91(1-2): 115–137.

Ekman, P. (1989). The argument and evidence about universals in facial expres-sions. Handbook of social psychophysiology: 143–164.

Ekman, P. and W. V. Friesen (1976). “Measuring facial movement.” Environmental psychology and nonverbal behavior 1(1): 56–75.

Ekman, P. and W. V. Friesen (1986). “A new pan-cultural facial expression of emotion.” Motivation and emotion 10(2): 159–168.

Ekman, P. and W. V. Friesen (1988). “Who knows what about contempt: A reply to Izard and Haynes.” Motivation and Emotion 12(1): 17–22.

Ekman, P., et al. (2013). Emotion in the human face: Guidelines for research and an integration of findings, Elsevier.

Ekman, P., et al. (2002). “Facial action coding system: The manual on CD ROM.” A Human Face, Salt Lake City: 77–254.

Ekman, P. and D. Keltner (1997). “Universal facial expressions of emotion.” Segerstrale U, P. Molnar P, eds. Nonverbal communication: Where nature meets culture: 27–46.

Freytag, A., et al. (2016). Chimpanzee faces in the wild: Log-euclidean cnns for predicting identities and attributes of primates. German Conference on Pattern Recognition, Springer.

Fridlund, A. J., et al. (1987). Facial expressions of emotion. Nonverbal behavior and communication, 2nd ed. Hillsdale, NJ, US, Lawrence Erlbaum Associates, Inc: 143–223.

Friesen, E. and P. Ekman (1978). “Facial action coding system: a technique for the measurement of facial movement.” Palo Alto 3.

Geisser, S. (1975). “The predictive sample reuse method with applications.” Journal of the American statistical Association 70(350): 320–328.

He, H. and E. A. Garcia (2009). “Learning from imbalanced data.” IEEE Transactions on knowledge and data engineering 21(9): 1263–1284.

Hinde, R. A. and T. Rowell (1962). Communication by postures and facial expressions in the rhesus monkey (Macaca mulatta). Proceedings of the Zoological Society of London, Wiley Online Library.

Jenny, A. B. and C. B. Saper (1987). “Organization of the facial nucleus and corticofacial projection in the monkey: a reconsideration of the upper motor neuron facial palsy.” Neurology 37(6): 930–930.

Krakauer, J. W., et al. (2017). “Neuroscience needs behavior: correcting a reductionist bias.” Neuron 93(3): 480–490.

Livneh, U., et al. (2012). “Self-monitoring of social facial expressions in the primate amygdala and cingulate cortex.” Proc Natl Acad Sci U S A.

Loos, A. and A. Ernst (2013). “An automated chimpanzee identification system using face detection and recognition.” EURASIP Journal on Image and Video Processing 2013(1): 49.

McLennan, K. M., et al. (2019). “Conceptual and methodological issues relating to pain assessment in mammals: The development and utilisation of pain facial expression scales.” Applied Animal Behaviour Science.

Mitz, A. R., et al. (2017). “Using pupil size and heart rate to infer affective states during behavioral neurophysiology and neuropsychology experiments.” Journal of Neuroscience Methods 279: 1–12.

Morecraft, R. J., et al. (2001). “Cortical innervation of the facial nucleus in the non-human primate: a new interpretation of the effects of stroke and related subtotal brain trauma on the muscles of facial expression.” Brain 124(1): 176–208.

Mosher, C. P., et al. (2016). “Tactile Stimulation of the Face and the Production of Facial Expressions Activate Neurons in the Primate Amygdala.” eNeuro 3(5).

Murphy, A. P. and D. A. Leopold (2019). “A parameterized digital 3D model of the Rhesus macaque face for investigating the visual processing of social cues.” Journal of Neuroscience Methods.

Padgett, C. and G. W. Cottrell (1997). Representing face images for emotion classification. Advances in neural information processing systems.

Panksepp, J. (2004). Affective Neuroscience: The Foundations of Human and Animal Emotions, Oxford University Press.

Pantic, M. and M. S. Bartlett (2007). Machine analysis of facial expressions. Face recognition, InTech.

Parr, L. A., et al. (2010). “Brief communication: MaqFACS: A muscle-based facial movement coding system for the rhesus macaque.” Am J Phys Anthropol 143(4): 625–630.

Pryluk, R., et al. (2019). “A Tradeoff in the Neural Code across Regions and Species.” Cell 176(3): 597–609.e518.

Pryluk, R., et al. (2020). “Shared yet dissociable neural codes across eye gaze, valence and expectation.” Nature 586(7827): 95–100.

Russell, W. M. S., et al. (1959). The principles of humane experimental technique, Methuen London.

Sariyanidi, E., et al. (2015). “Automatic Analysis of Facial Affect: A Survey of Registration, Representation, and Recognition.” IEEE TRANSACTIONS ON PATTERN ANALYSIS AND MACHINE INTELLIGENCE 37(6): 1113–1133.

Tukey, J. (1958). “Bias and confidence in not quite large samples.” Ann. Math. Statist. 29: 614.

Vick, S.-J., et al. (2007). “A Cross-species Comparison of Facial Morphology and Movement in Humans and Chimpanzees Using the Facial Action Coding System (FACS).” Journal of Nonverbal Behavior 31(1): 1–20.

Waller, B., et al. (2020). “Measuring the evolution of facial ‘expression’using multi-species FACS.” Neuroscience & Biobehavioral Reviews.

Welt, C. and J. H. Abbs (1990). “Musculotopic organization of the facial motor nucleus in Macaca fascicularis: a morphometric and retrograde tracing study with cholera toxin B-HRP.” Journal of Comparative Neurology 291(4): 621–636.

Witham, C. L. (2017). “Automated face recognition of rhesus macaques.” Journal of Neuroscience Methods.

